# Towards a mechanistic understanding of the role of error monitoring and memory in social anxiety

**DOI:** 10.1101/2023.09.14.557662

**Authors:** Kianoosh Hosseini, Jeremy W. Pettit, Fabian A. Soto, Aaron T. Mattfeld, George A. Buzzell

## Abstract

Cognitive models state social anxiety (SA) involves biased cognitive processing that impacts what is learned and remembered within social situations, leading to the maintenance of SA. Neuroscience work links SA to enhanced error monitoring, reflected in error-related neural responses arising from mediofrontal cortex (MFC). Yet, the role of error monitoring in SA remains unclear, as it is unknown whether error monitoring can drive changes in memory, biasing what is learned or remembered about social situations. Thus, we developed a novel paradigm to investigate the role of error-related MFC theta oscillations (associated with error monitoring) and memory biases in SA. EEG was collected while participants completed a novel Face-Flanker task, involving presentation of task-unrelated, trial-unique faces behind target/flanker arrows on each trial. A subsequent incidental memory assessment evaluated memory biases for error events. Severity of SA symptoms were associated with greater error-related theta synchrony over MFC, as well as between MFC and sensory cortex. SA was positively associated with memory biases for error events. Consistent with a mechanistic role in biased cognitive processing, greater error-related MFC-sensory theta synchrony during the Face-Flanker predicted subsequent memory biases for error events. Our findings suggest high SA individuals exhibit memory biases for error events, and that this behavioral phenomenon may be driven by error-related MFC-sensory theta synchrony associated with error monitoring. Moreover, results demonstrate the potential of a novel paradigm to elucidate mechanisms underlying relations between error monitoring and SA.

## Introduction

Social anxiety (SA) is characterized by an extreme, persistent fear of social situations [1] and is one of the most pervasive, chronic, and difficult-to-treat anxiety disorders [2]. Cognitive models describe how SA symptoms are maintained or worsened over time [3], [4], informing efficacious treatment approaches [5]. Nonetheless, treatment outcomes remain suboptimal [6]. Clinical neuroscience has identified neural markers of risk for SA [7]–[9], yet it is unclear how supplanting psychological/cognitive measures with neural markers will translate into improved treatment. Thus, there is a need to move beyond “neural markers” towards identification of *neural mechanisms* implicated in SA that can be targeted/manipulated in treatment. Toward these ends, the current proof-of-concept study draws on cognitive models of SA and emerging clinical neuroscience research to investigate the role of error-related neural oscillations and memory biases in SA.

Cognitive models state that the maintenance of SA is driven by biased cognitive processing, which impacts what is learned and remembered within social situations [3], [4]. Specifically, cognitive models state that fear and worry about social situations leads to greater self-focus, self-monitoring, and attention toward negative aspects of performance (e.g., making mistakes/errors). As a result, negative aspects of performance become more salient and thus better encoded/remembered, negatively biasing self-assessments (i.e., post-event processing) and maintaining SA [3], [4]. Empirical work supports these assertions: SA and self-focus predict memory biases for negative aspects of performance and worse self-evaluations following social situations, ultimately maintaining or worsening SA [10]–[16]. However, work is needed to bridge these cognitive models with emerging findings from neuroscience to elucidate neural mechanism(s) implicated in SA [17].

Within cognitive neuroscience, “error monitoring” refers to the process of self-monitoring and detecting one’s mistakes, associated with neural activity arising from medial frontal cortex (MFC) [18]–[21]. EEG is particularly well-suited for studying error monitoring, given high temporal resolution [22] and sensitivity to oscillatory patterns (power/phase relations; [23]). Most work focuses on two related EEG measures: the Error-Related Negativity (ERN) [24], [25] and error-related MFC theta oscillations [18], [26], both recorded over MFC and localized, at least in part, to neural sources within MFC [27]–[30]. Theta oscillations exhibit maximal increases in power (magnitude) and synchrony (phase alignment) over MFC following error responses [18], [29], [31], [32]. Moreover, error responses elicit enhanced theta synchronization between MFC and task-relevant brain regions [31], [33], [34], in line with the MFC reflecting a central node in an extended network that detects the need for and recruits control following errors [35], [36].

Extensive work demonstrates EEG-based measures of error monitoring—recorded over MFC—are linked to anxiety [37]–[40], including SA [41]–[45]. For high SA individuals, error monitoring is particularly increased within social situations [46]–[48]. However, the directionality of associations between error monitoring and (social) anxiety remain unresolved. One class of theories suggests enhanced error monitoring is a symptom of anxiety and does not play a causal role, such that error-monitoring reflects compensatory efforts to control behavior due to distracting effects of anxiety [39], [49]. Another class of theories suggests error monitoring predicts “risk” for anxiety, in part based on their prospective relations (e.g., [44], [50]–[52]), but does not specify whether error monitoring plays a causal role [38], [40]. Critically, for error monitoring to play a causal role in social anxiety, it must impact learning/memory to drive lasting changes in SA. Otherwise, any effects of error monitoring on cognition and behavior would be transitory in nature.

As previously described, cognitive models of SA state that self-focus, self-monitoring, and attention to negative aspects of performance increase error salience and subsequent encoding, biasing self-evaluations (post-event processing) and maintaining SA [3], [4]. As a neural extension of these models, we propose that error-related MFC theta oscillations provide a neural mechanism for error monitoring to increase the likelihood of encoding error events, which could contribute to the maintenance or exacerbation of SA. First, SA is associated with increased error monitoring [41], [46], [48], [53] as well memory biases for negative aspects of performance (e.g., errors; [10]– [13], [15], [16]. Second, MFC theta oscillations are not only associated with error monitoring [31], [54], but separate research demonstrates fundamental associations with memory: MFC theta oscillations during encoding predict increased likelihood of later recall [55]–[58]. Third, error monitoring is also known to drive increases in attention [33], [34], [59], attention is known to rely on theta band phase synchronization [60]–[63], and the role of attention in memory encoding is well established [64]–[66]. Collectively, we propose that error-related MFC theta oscillations (associated with error monitoring) may increase the likelihood that error events are encoded and later remembered. Further, we anticipate that error-related increases in theta phase synchronization (connectivity) between MFC and task-relevant sensory cortices (e.g., occipital-parietal for visual information) associated with attention to (and thus, encoding of) error-related information may be particularly relevant.

Prior studies of memory biases in SA typically assess memory following dynamic social interactions (e.g., giving a speech; [11]–[13], [15], [16]). However, the less-structured nature of these approaches limits neural assessments of error monitoring. Similarly, work linking SA to heightened error monitoring typically employs computer tasks (e.g., Flanker) using limited stimulus sets [67], which constrains attempts to probe memory of individuals’ error events, given the lack of trial-unique contexts. Addressing these limitations, we created a novel paradigm involving presentation of trial-unique faces during a flanker task—performed under social observation—followed by incidental memory assessment. In this proof-of-concept study, use of a flanker task allowed for extracting known patterns of error-related MFC theta oscillations: power/phase over MFC and phase relations between MFC and task-relevant sensory (visual) cortices. Employing a subsequent incidental memory assessment and comparing recognition memory for faces previously presented during error (vs. correct) events, allowed for indexing memory biases for error events. Leveraging these measures, we tested three hypotheses regarding the role of error monitoring and memory biases in SA: 1) SA is associated with memory biases for error events; 2) SA is associated with increased error monitoring, reflected in enhanced error-related MFC theta oscillations; and 3) error-related MFC theta oscillations, at the time of encoding, are associated with subsequent memory biases for error events.

## Methods

### Participants

Fifty-four healthy adult individuals (M = 23.48 years, SD = 3.45; 48 f, 6 m) provided informed consent prior to participation and received either monetary compensation or course credit. All participants were fluent in English and had no prior head injury causing loss of consciousness.

Given that this was a proof-of-concept study, a relatively shortened experimental task was employed; this resulted in the exclusion of 20 participants that committed fewer than 8 errors (insufficient data for error-related analyses). Two more participants were excluded due to experimental inconsistencies/error, resulting in a total of 32 participants (M = 23.5 years, SD = 3.31; 29 f, 3 m) for behavioral analyses. Of these 32 individuals, seven did not have EEG recorded, and one was excluded from EEG analyses due to having an insufficient number of artifact-free trials per condition of interest, in line with prior work: [68], [69]. Thus, 24 participants (M = 23.68 years, SD = 3.68; 22 f, 2 m) were included in EEG data analyses. Note the sample imbalance for biological sex results from unbiased enrollment of participants from a predominately female undergraduate psychology student population [70].

### Assessment of SA symptoms

Participants self-reported on their SA symptom levels via the 7-item Social Anxiety scale derived from the Screen for Adult Anxiety Related Disorders (SCAARED) [71]. Questionnaire items are presented on a 3-point Likert scale (0 = not true or hardly ever true, 1 = somewhat true or sometimes true, 2 = very true or often true) and a higher score represents more severe symptoms of SA. Cronbach’s α for the SCAARED Social Anxiety scale was 0.79 in this study.

### Procedure

To investigate the role of error monitoring on subsequent memory, participants first completed a novel Face-Flanker task while EEG was recorded. To create a context of social evaluation, participants were explicitly told their performance would be monitored and evaluated while they completed the Face-Flanker task. Subsequently, participants performed a surprise incidental memory assessment, in which all faces (n = 160) from the Face-Flanker task along with 80 never-before-seen faces were presented. Participants also performed a facial expression encoding task, which is not discussed further, as it is beyond the scope of the current report.

### Face-Flanker task

Participants completed a modified Flanker task (Figure 1); on each trial, participants were presented with an array of five arrows, with a trial-unique neutral face image from the Chicago face database in the background [72], [73]. Participants used their right/left thumbs to indicate the (right/left) direction of a target arrow via button press. Flanking arrows were oriented in the same (congruent) or opposite (incongruent) direction as the target arrow. Flanking arrows always appeared first and remained on the screen for 150 ms prior to the target arrow appearing; all arrows then remained on the screen for 200 ms prior to disappearing synchronously. A fixation rectangle was maintained on-screen throughout each block, positioned in the center of the screen, and surrounding the arrow array. The background face was maintained onscreen for the duration of the trial, which was randomly jittered between 3500-4000 ms. Stimuli were presented on a 15-inch Lenovo Legion 7i laptop running Windows 10 with PsychoPy version 2021.2.3 [74]. Participant responses were recorded throughout the duration of each trial via the Black Box ToolKit (BBTK) response pad (The Black Box ToolKit Ltd., Sheffield, UK). To ensure attentiveness of participants, trials (M = 0.04, SD = 0.19) with a reaction time (RT) faster than 150 ms were removed from further analyses.

**Figure 1.** The Face Flanker Task. A single trial from the Face Flanker task is shown, to include the stimulus onset asynchrony between the target and flanker arrows. A trial-unique background face from the Chicago face database and the fixation rectangle are shown. [*This figure was removed per bioRxiv policy to remove images with faces prior to posting. Note, figure 1 is available from the authors upon request; Faces are drawn from the Chicago face database [72], [73]]*

Participants completed 5 blocks of 32 trials (160 trials total), with an equal mix of congruent/incongruent trials in each block. To facilitate adequate error rates, feedback was displayed following each block [75]. If accuracy was above 75% but below 90%, *“Good job”* was displayed; “Respond *faster”* or *“Respond more accurately” were* presented when the accuracy was above 90% or below 75%, respectively. At task completion, participants self-reported the number of errors made.

### Incidental memory assessment

Following completion of the Face-Flanker task, participants completed an incidental, self-paced memory assessment. Participants indicated whether they recognized each individually presented face as “new” or “old” (recognized as previously appearing during the Face-Flanker task). This task consisted of 240 trials, with 160 old faces drawn from the Face-Flanker task and 80 new (foil) faces randomly intermixed. Faces were presented across a total of two blocks (120 faces/block). Each face was presented until a button press response was made using left/right thumbs; response mappings between left/right thumbs and new/old responses were counterbalanced across participants. To ensure attentiveness of participants during the task, trials (M = 0.47, SD = 0.80) with a reaction time (RT) faster than 200 ms were removed; all participants had less than 20% of trials removed. The same computer equipment, software, and peripherals employed in the Face-Flanker task were used.

### Memory bias for error events

To index memory bias for error events, we evaluated the degree to which participants varied in recognition memory performance for face images that originally appeared during error events (error trials from the Face-Flanker task) relative to correct events. We first computed separate hit rate scores (% correctly identified) for faces that originally appeared on error and correct Face-Flanker trials (See equations 1 and 2). We then computed a difference score to index memory bias for error events by subtracting hit rates for correct events from hit rates for error events (equation 3). This hit rate difference score was used in subsequent statistical analyses (see preliminary analyses section). Note that for all analyses, only faces that appeared on incongruent (error/correct) trials were analyzed to obviate a confound of stimulus congruency and isolate error-related effects of interest (errors are more common on incongruent trials) [31], [67]; throughout the manuscript, we refer to incongruent-error and incongruent-correct trials as “error” and “correct” for simplicity.

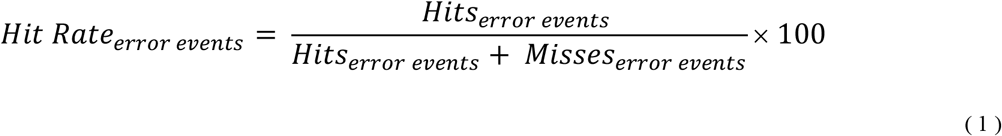

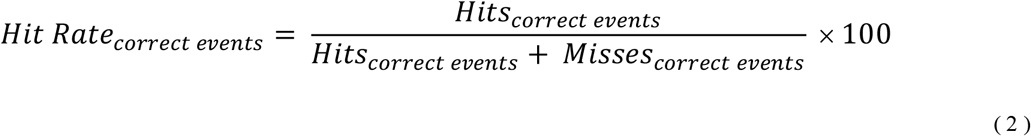

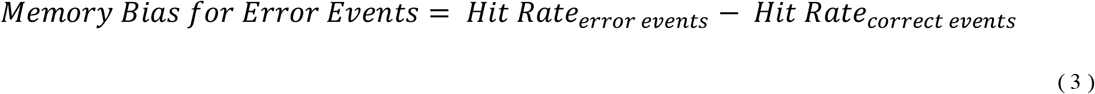

### EEG acquisition and preprocessing

To assess error monitoring during the face-flanker task, 64-channel EEG was collected via a Brain Products actiCHamp amplifier and BrainVision Recorder software (Brain Products GmbH, Munich, Germany) at 1000 Hz. A 64-channel EasyCap custom EEG cap (EasyCap GmbH, Herrsching, Germany) was used (see Figure 2 for cap layout). Impedance was reduced to a targeted level of ≤ 25 kΩ prior to data collection. EEG electrodes were referenced to electrode 1 (∼FCz; see figure 2) during recording and re-referenced to the average of all electrodes during preprocessing. EEG data were preprocessed using MATLAB R2021b (MathWorks Inc., Sherborn, MA, USA), the EEGLAB toolbox, and a modified version of the MADE pipeline [76], [77]. As part of preprocessing, data were segmented into 3-second epochs (-1 to 2 seconds, relative to the response). Complete details of the EEG preprocessing stream can be found in the supplement.

**Figure 2.**
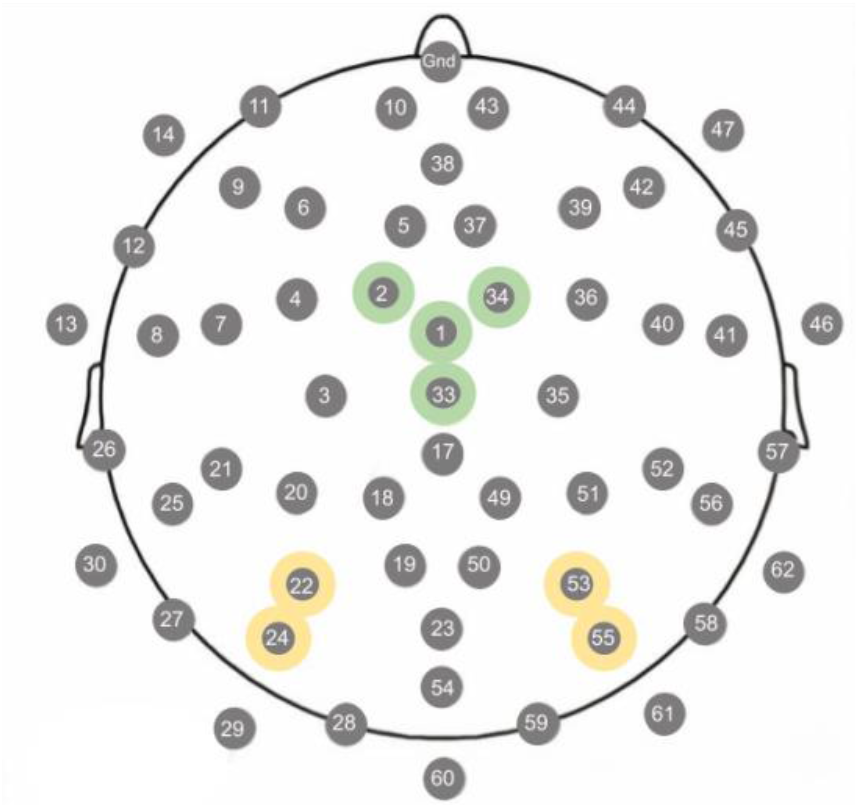
64-channel EasyCap EEG cap layout. The selected electrodes over the MFC and bilateral sensory (visual) regions are shown in green and yellow circles, respectively. Their polar coordinates (theta, phi) and closest equivalent electrode on the standard 10-5 localization system are as follows: E1 (17, 90) FCz; E2 (-34, -60) FFC1h; E33 (0, 0) Cz; E34 (34, 60) FFC2h; E22 (-68, 54) PPO5h; E24 (-85, 56) PO7; E53 (68, -54) PPO6h; E55 (85, -56) PO8.

### Error-Related MFC Theta Oscillations

MATLAB scripts (based on work by: [23], [78]) were used to compute response-locked time-frequency (TF) power and phase relations—within and between channels—for epochs of interest. Mirroring the behavioral analyses, all EEG analyses focused on incongruent error/correct trials to obviate a confound of stimulus congruency and isolate error-related effects of interest [31], [67]. To increase computational efficiency, EEG data were downsampled to 250 Hz following decomposition of TF activity. All TF measures were normalized relative to a -400 to -200 pre-response baseline.

#### MFC theta power

Morlet wavelets were convolved with each epoch to estimate spectral power (oscillation magnitude) between 1-30 Hz, divided into 59 logarithmically-spaced steps [23]. The number of wavelet cycles increased from 3 (at 1 Hz) to 10 (at 30 Hz) to balance time/frequency precision [23], [78]. TF power was separately computed for each epoch of interest, for all channels, before averaging across epochs within a given condition to calculate total power. In line with prior studies [18], [79], for each condition of interest, mean response-locked MFC theta power was analyzed within a region of interest (ROI) spanning 4-7 Hz and the first 250 ms following response within a cluster of electrodes located over MFC (∼FCz and surrounding electrodes: 1, 2, 33, and 34; see Figure 2).

#### MFC theta intertrial phase clustering

Intertrial Phase Clustering (IPC) reflects the consistency of oscillatory phase angles across trials for a given frequency/timepoint, relative to an event of interest (in this case, error and correct responses) [23]. IPC is scaled between 0 and 1, with 0 denoting random phase alignment and 1 denoting perfect phase alignment. To compute IPC for a given TF point, the phase angle difference across trials was taken before averaging. In line with prior studies [18], [79], for each condition of interest, MFC theta IPC was analyzed using the same TF ROI (4-7 Hz, 0-250 ms) and MFC electrode cluster defined above. To avoid biases associated with calculating phase-based measures using unequal trials counts, a subsampling procedure was implemented [23], [78]: six trials were randomly selected per condition and the subsampling process was repeated 100 times before averaging.

#### MFC-sensory theta weighted phase-lag index

Weighted Phase-lag Index (wPLI) reflects the consistency of phase angles between channels (connectivity), across trials, for a given TF point [23], [78]. Of note, wPLI minimizes effects of volume conduction by de-weighting phase angle differences near-zero, allowing for the analyses in raw channel-space [80]. To compute wPLI for a given TF point, the phase angle differences between channels was taken, and the sign of the imaginary part of the cross-spectral density for each electrode pair over trials was averaged. Based on our hypothesis that enhanced error-related MFC theta oscillations (central to error monitoring) are associated with attention towards error events (increasing the likelihood of encoding), and given the visual nature of our task, we computed wPLI between a seed electrode centered within the MFC cluster defined above (∼FCz: electrode 1; see Figure 2) and a bi-lateral cluster of electrodes (∼PO7/PO8: electrodes 22, 24, 53, 55; see Figure 2) located over visual sensory regions (occipital-parietal cortex; [81], [82]. For each condition of interest, MFC-sensory theta wPLI was analyzed using the same TF ROI (4-7 Hz, 0-250 ms) and subsampling procedure (6 trials, 100 repetitions) described above.

### Analytic Plan

All statistical analyses were conducted using R [83]. One-tailed statistical tests were used for directional hypotheses; two-tailed tests were otherwise employed. Where appropriate, control over the family-wise error rate was achieved via a Holm-ŠÍdák correction; in such cases, we report both uncorrected and corrected p-values.

#### Preliminary Analyses

To confirm the presence of standard congruency effects [67] during the Face-Flanker task, two non-parametric paired-sample one-tailed Wilcoxon tests were performed to compare accuracy rates between incongruent and congruent trials, as well as to compare mean RT between correct incongruent and congruent trials. To confirm the presence of error-related changes in MFC theta oscillations (an index of error monitoring) during the Face-Flanker task, an *a priori* series of paired-sample one-tailed t-tests were employed to compare error and correct trial responses for MFC theta power, MFC theta IPC, and MFC-sensory theta wPLI; correction for multiple comparisons was applied to this family of tests. To assess overall recognition memory performance for faces originally presented during error vs. correct events, a paired-sample two-tailed t-test was used to compare their respective hit rates.

Following these preliminary analyses, error-related difference scores (error-correct) were computed for MFC theta power, MFC theta IPC, and MFC-sensory theta wPLI to carry out a series of analyses testing our central hypotheses. As previously described, we also computed a difference score to index memory bias for error events by subtracting hit rates for correct events from error events (equation 3).

#### Statistical Analyses

To test whether higher SA symptom levels were associated with memory bias for error events, we carried out an *a priori* one-tailed Pearson correlation test of whether SCAARED-Social scores were significantly correlated with memory bias for error events difference scores.

Next, to confirm that higher SA symptom levels were associated with error-related MFC theta oscillations at the time of encoding—during the Face-Flanker task—we carried out an *a priori* series of one-tailed Pearson correlation tests between SCAARED-Social scores and error-related differences scores for: MFC theta power, MFC theta IPC, and MFC-sensory theta wPLI. Correction for multiple comparisons was applied to this family of tests.

After determining which error-related MFC theta oscillations measure(s) were significantly correlated with SA symptom levels, we further tested whether these same error-related MFC theta oscillations measure(s) predicted memory bias for error events difference scores via a series of regression analyses (one-tailed tests); correction for multiple comparisons was again applied to this family of tests. We also carried out a control analysis to rule out the possibility that memory biases for error events were instead driven by stimulus-evoked responses to face onsets (see supplement).

## Results

### Preliminary behavioral results

Consistent with prior Flanker task studies [67] participants responded less accurately on incongruent (Md = 81.87%, n = 32) compared to congruent trials (Md = 97.50 %, n = 32) trials, *z* = -5.063, *p* < 0.001, Cohen’s d = 2.719. Similarly, participants responded more slowly on incongruent-correct (Md = 557.07 ms, n = 32) compared to congruent-correct (Md = 487.48 ms, n = 32) trials, *z* = -6.338, *p* < 0.001, Cohen’s d= 1.331.

The average hit rate of participants in the surprise incidental memory assessment was 46.37% (SD = 13.40%), consistent with studies evaluating memory performance for task-irrelevant stimuli using a comparable number of images [84]. On average, participants did not differ in terms of recognizing faces originally presented during error vs. correct events, *t*(31) = 0.592, *p* = 0.558, Cohen’s d = 0.088.

### SA symptoms positively relate to memory biases for error events

Consistent with our hypotheses, SA symptom levels (assessed via SCAARED-social) were positively associated with memory biases for error events (better recognition of faces that previously appeared during error vs. correct events), *r*(30) = 0.451, *p* = 0.005.

**Figure 3.**
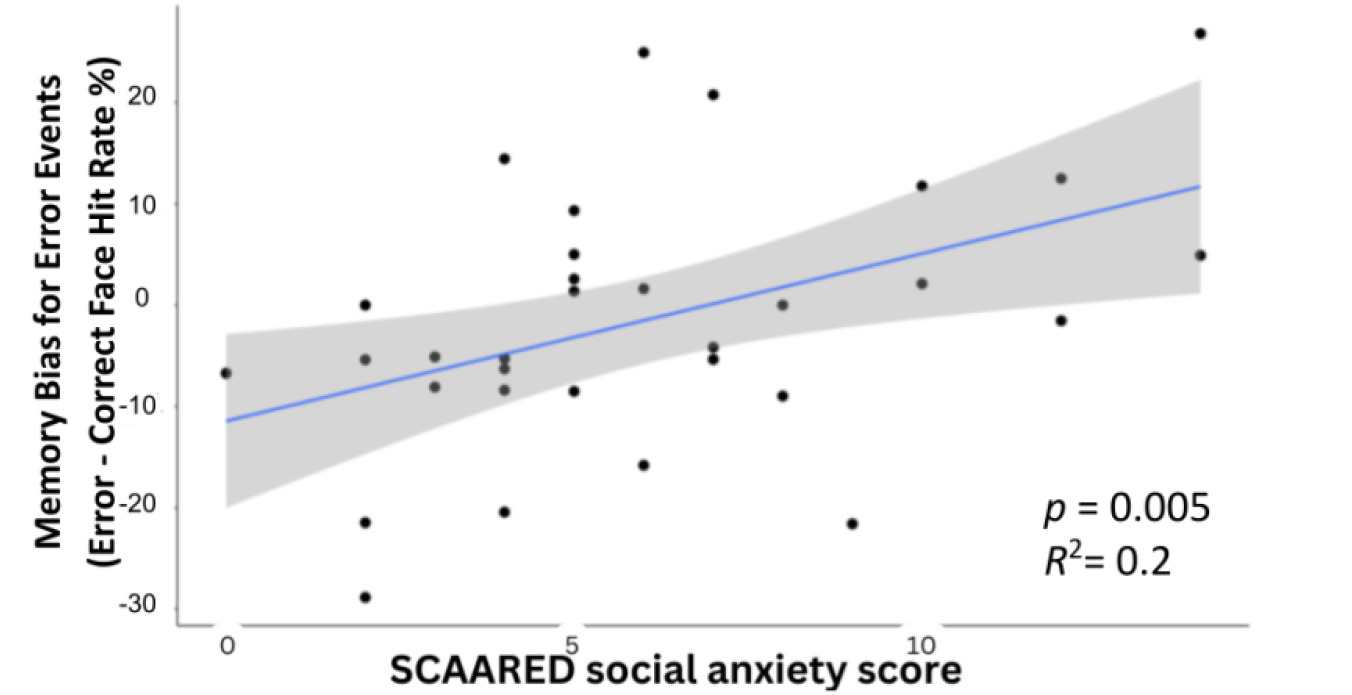
The relationship between SCAARED-social scores and memory bias for error events. SCAARED-social scores were significantly associated with memory biases for error events (better recognition memory performance for faces that originally appeared during error vs. correct events).

### Error-Related MFC Theta Oscillations During the Face Flanker Task

To assess error-related MFC theta oscillations (associated with error monitoring) during the Face-Flanker task, we first performed a series of a preliminary analyses comparing response-locked theta oscillations for error vs. correct trials. In line with prior error monitoring work, error responses (relative to correct) were associated with a robust increase in MFC theta power, *t*(23) = 8.775, *p* < 0.001 (*p*_adj_ < 0.001), Cohen’s d = 1.79. Similarly, MFC theta IPC significantly increased for error (vs. correct) responses, *t*(23) = 1.87, *p* = 0.037 (*p*_adj_ = 0.037), Cohen’s d = 0.38. Error (vs. correct) responses were also associated with a significant increase in MFC-sensory theta wPLI, t(23) = 3.15, *p* = 0.002 (*p*_adj_ = 0.004), Cohen’s d = 0.64. This latter result is consistent with the notion that error monitoring involves rapid engagement of visual sensory regions, which could in turn impact the encoding contextual information during an error event. See Figure 4 for a depiction of these results.

**Figure 4:**
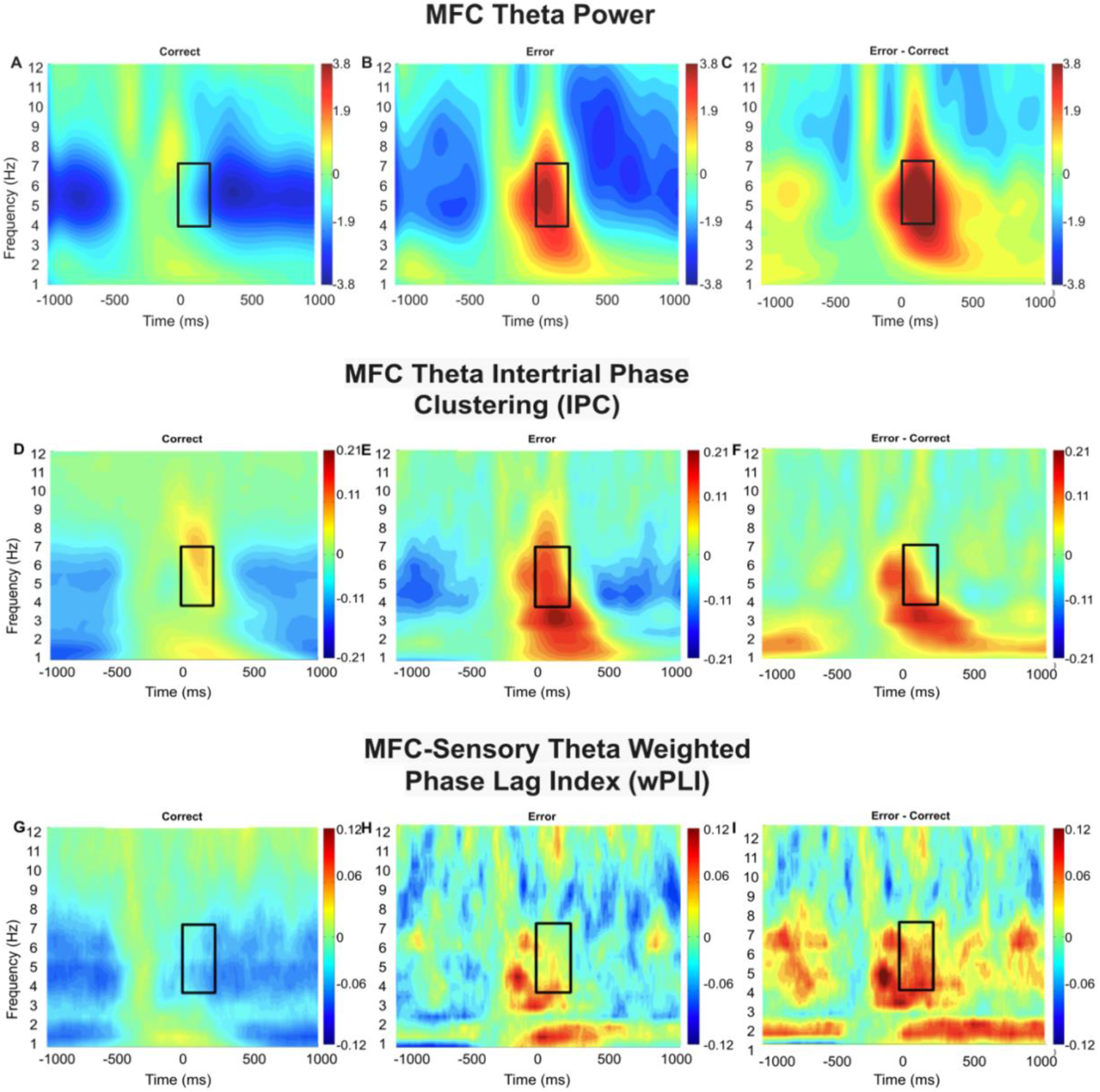
Error-related MFC theta oscillations. In all plots, 0 ms corresponds to the time of response; black-box overlays depict the a priori time-frequency (TF) region of interest used for analysis (4-7 Hz, 0-250 ms). All plots and analyses employ incongruent error/correct trials only to avoid stimulus-related confounds (see text). (A, B, C) MFC theta power TF plots for correct, error, and the error – correct difference, respectively; (D, E, F) MFC theta IPC TF plots for correct, error, and the error – correct difference, respectively; (G, H, I) MFC-Sensory theta wPLI TF plots for correct, error, and the error – correct difference, respectively.

### SA Symptoms Positively Relate to Error-Related MFC Theta Oscillations

To test whether error-related MFC theta oscillations (associated with error monitoring) were more pronounced for individuals higher in SA symptom levels, we tested whether SCAARED-social scores correlated with error-correct difference scores for each of the error-related MFC theta measures described above (MFC theta power, MFC theta IPC, MFC-sensory theta wPLI). Whereas SCAARED-social scores did not significantly relate to error-related MFC theta power, *r*(22) = 0.254, *p* = 0.115 (*p*_adj_ = 0.115), SCAARED-social scores were significantly associated with error-related MFC theta IPC, *r*(22) = 0.403, *p* = 0.025 (*p*_adj_ = 0.0496). Similarly, SCAARED-social scores were significantly related to MFC-sensory theta wPLI: higher SA symptom levels were positively associated with error-related MFC-sensory theta wPLI, *r*(22) = 0.469, *p* = 0.010 (*p*_adj_ = 0.030). These results are consistent with the notion that error-related MFC theta oscillations (associated with error monitoring) are enhanced in individuals high in SA. See Figure 5 for a depiction of these results.

**Figure 5:**
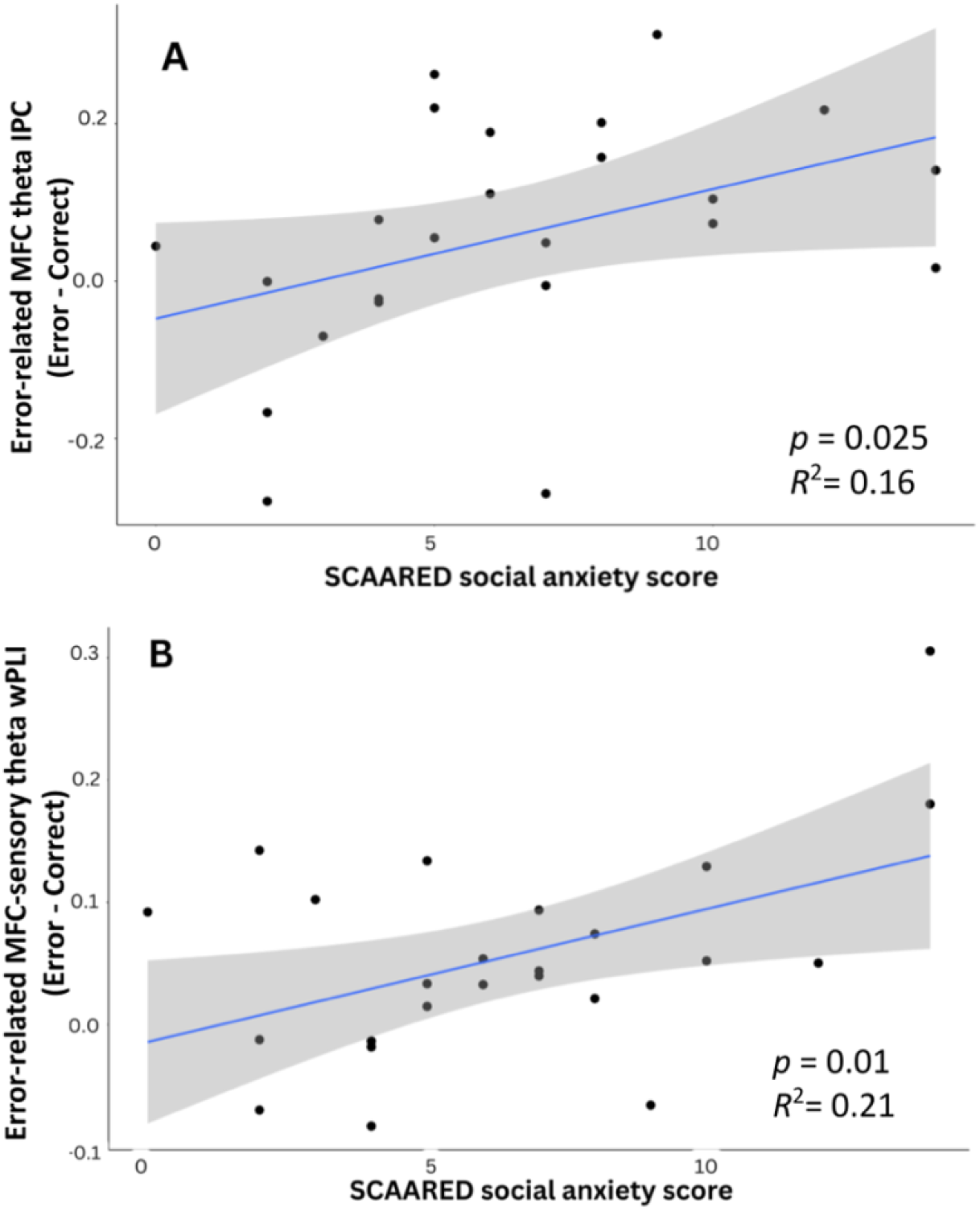
Associations between SA symptoms and MFC theta oscillations. Higher SA symptom levels are positively associated with: (A) error-related MFC theta IPC and (B) error-related MFC-sensory theta wPLI.

### Error-related MFC theta oscillations predict subsequent memory biases for error events

Given that higher SA symptom levels positively related to both error-related MFC theta IPC and MFC-Sensory theta wPLI, we tested whether either of these error-related MFC theta measures also predicted subsequent memory biases for error events. Error-related MFC-Sensory theta wPLI related positively to memory bias for error events difference scores (better recognition of faces that previously appeared during error vs. correct events), β = .428, *t*(1,22) = 2.218, *p* = 0.019 (*p*_adj_ = 0.037). Error-related MFC IPC did not exhibit similar relations with memory bias for error events difference scores, β = 0.067, *t*(1, 22) = 0.316, *p* = 0.377 (*p*_adj_ = 0.377). These data are consistent with the hypothesis that error monitoring drives memory biases for error events: heightened error-related engagement between MFC and visual sensory regions may drive enhanced encoding of error-related contextual information present at the time an error is committed. Further supporting this interpretation, a supplemental analysis ruled out the possibility that memory biases for error events could have been driven by stimulus-evoked neural responses to face onsets (see supplement).

**Figure 6:**
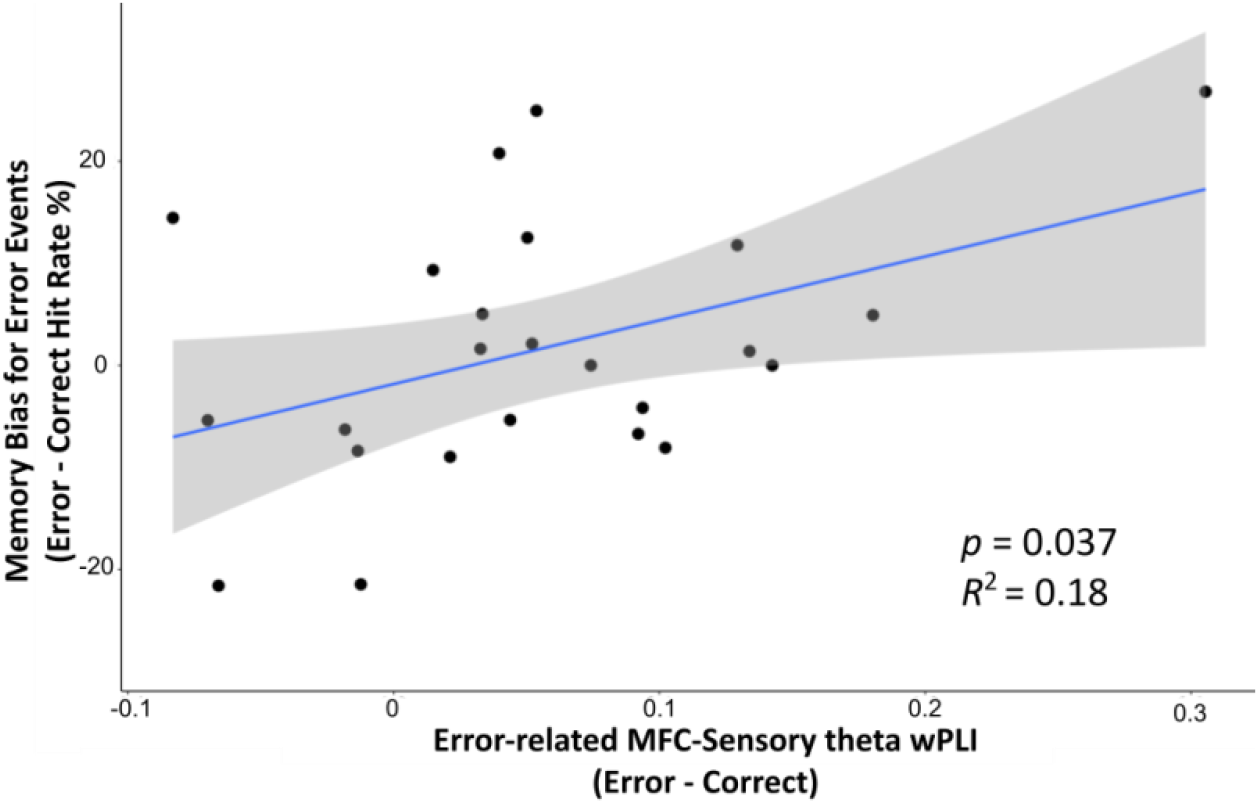
The relationship between Error-related MFC-sensory theta wPLI and memory bias for error events. Error-related MFC-sensory theta wPLI is significantly associated with memory bias for error events (better recognition of faces that previously appeared during error vs. correct events).

## Discussion

Bridging cognitive models of SA with recent neuroscience findings, the current study investigated the putative role of error-related MFC theta oscillations (associated with error monitoring) and memory biases in SA. Participants completed the novel Face-Flanker task, allowing measurement of error monitoring, followed by an incidental memory assessment, providing an index of memory biases for error events (degree to which error vs. correct events from the Face-Flanker were preferentially remembered). SA symptoms were positively associated with memory biases for error events. Within the same paradigm, SA symptoms were also positively associated with error-related MFC theta oscillations at the time of encoding. Specifically, SA was associated with enhanced error-related MFC theta IPC (synchrony over MFC), as well as enhanced error-related MFC-sensory theta wPLI (synchrony between electrode sites located over MFC and visual-sensory cortex). Additionally, error-related MFC-sensory theta wPLI—at the time of encoding—further predicted subsequent memory biases for error events. Collectively, these findings provide proof-of-concept support for a neural mechanism implicated in SA: memory biases following social situations may arise, in part, from enhanced error-related MFC theta oscillations that increase the likelihood that error events are encoded and subsequently remembered. Future work should seek to replicate and extend these findings, leveraging the Face Flanker task in combination with longitudinal assessment of state/trait SA symptoms to directly test whether the proposed neural mechanism is causally implicated in the maintenance or worsening of SA.

### SA associated with memory biases for error events

Consistent with prior work using less-structured paradigms (e.g., memory assessments following a speech or other social interaction; [11]–[13], [15], [16], our behavioral data suggest error events are better remembered for individuals high in SA. While we interpret such memory biases as arising from enhanced error monitoring in high SA individuals, this cannot be confirmed based on behavioral data alone. This is because alternatively, high SA individuals could simply be more distracted by faces to begin with (preferentially attending to and encoding faces), which then causes errors to occur, as opposed to error monitoring driving the encoding of error events. However, our neural data present a pattern of results consistent with our hypothesis that memory biases for error events are driven by heightened error monitoring. SA symptoms were positively associated with heightened memory biases for error events as well as heightened error-related MFC theta oscillation patterns indicative of enhanced error monitoring. In particular, high SA individuals exhibited enhanced error-related MFC-sensory theta wPLI, which further predicted subsequent memory biases for error events. Moreover, we identified no evidence in favor of the alternative interpretation, as stimulus-evoked neural responses to face onsets were not associated with SA nor subsequent memory biases. Collectively, these data not only demonstrate that SA is associated with memory biases for error events, but also provide evidence that such memory biases may arise as the result of error monitoring (error-related MFC theta oscillations).

### SA associated with enhanced error monitoring

The observed relations between SA and enhanced error-related MFC theta oscillations are consistent with prior work linking SA to error monitoring [41]–[48]. It is worth noting that the majority of prior work investigating relations between (social) anxiety and error monitoring has focused on time-domain (ERP) analyses of the ERN. However, given prior work linking MFC theta to both error monitoring and memory [26], [31], [55]–[58], we chose to employ a TF-analytic approach and focus on error-related MFC theta oscillations in the current report. We found that SA was associated with synchrony-based theta measures (IPC and wPLI), but not theta power. This link between SA and synchrony-based measures of error-related MFC theta is noteworthy, given that theta synchrony, as opposed to theta power, has also been shown to be more closely related to the ERN [29], [85], [86]. Thus, our findings are consistent with prior work demonstrating that SA is associated with an enhanced error monitoring, as measured by the ERN [41], [42].

### Neural mechanism underlying the link between error monitoring and memory biases

Current theoretical models of the link between error monitoring and anxiety propose that error monitoring either reflects a downstream symptom of anxiety [39], [49], or that error monitoring predicts “risk” for anxiety without specifying whether error monitoring plays a causal role [38], [40]. However, if error monitoring is to instead play a causal role in the etiology of SA, this requires a mechanism by which error monitoring could impact learning/memory to produce lasting changes in cognition and behavior. Our finding that error monitoring predicts memory biases for errors introduces the possibility that error monitoring may play a causal role in SA. As previously described, cognitive models of SA state that self-focus, self-monitoring, and attention to negative aspects of performance increase error salience and subsequent encoding, ultimately biasing self-evaluations (post-event processing) and maintaining SA [3], [4]. As a neural extension of these models, it is possible that increased error monitoring directly contributes to SA by impacting memory, biasing self-evaluations (post-event processing) and maintaining SA. The current study provides support for the link between error monitoring and memory, identifying error-related MFC theta oscillations as a neural mechanism by which error monitoring may increase the likelihood of encoding error events. Moreover, we demonstrate that SA symptoms are positively associated with enhanced error monitoring as well as memory biases for error events. The next logical step is to replicate and extend these findings, to test if memory biases for error events, driven by error monitoring, mediate longitudinal changes in state/trait SA. Similarly, associations with post-event processing [12], [15], [87] should be studied. For example, one possibility is that error monitoring drives memory biases for error events, which then skew post-event processing towards recollection of more negative aspects of behavior. Alternatively, post-event processing might interact with error monitoring to predict the degree to which memory biases for error events are maintained over time. Either of these possibilities could lead to the maintenance or worsening of SA.

### Broader implications of the identified link between error monitoring and memory

It is worth noting that observed relations between error-related MFC-sensory theta wPLI and memory biases for error events were present for all participants, regardless of SA symptoms. That is, although participants in our study did not exhibit memory biases for error events at the behavioral level, on average, we did find that individual variation in error-related MFC-sensory theta wPLI was predictive of individual variation in memory biases for error events. In other words, individuals that exhibited the strongest neural responses, at the time of encoding, were most likely to exhibit later memory biases for error events. Other recent behavioral work has found that, within the general population, either error events [84] or post-error events [88] are better remembered. Thus, although we did not find evidence for such behavioral effects, on average, it is possible that such effects could be detected within a larger sample. Regardless, our data provides the first evidence that individual variation in error-related MFC theta oscillations predict the degree to which error events are remembered. These data point to a potential neural mechanism underlying memory biases for error events that should be investigated in larger studies, not only in relation to social anxiety, but also within the general population.

It is also worth noting that another recent study did not identify a significant relation between error-related MFC theta oscillations and memory (in the general population; [89]. However, our data suggests two possible reasons for this difference across studies. First, the study by Zheng and wynn [89] only investigated relations between error-related MFC theta power (not synchrony) and memory. Our study found that a synchrony-based measure (wPLI) was associated with memory biases for error events, thus, theta synchrony may be more closely tied to the likelihood that error events are committed to memory. Second, whereas the study by Zheng and wynn [89] assessed memory by asking participants to recall the number of errors they made, in the aggregate, we indexed memory for error events by assessing recognition of images present on error trials. Thus, it is possible that these approaches rely on different forms of memory [90], [91] and/or differ in the resolution of memory assessment they provide (i.e., assessment of individual error events vs. aggregate estimates). Given that this is the first study to identify a link between error-related MFC theta oscillations and memory biases for error events, further work is needed to replicate and extend these findings, providing a more detailed characterization of the link between error-related MFC theta oscillations and memory.

### Limitations and future directions

The current report introduces a novel paradigm and presents proof-of-concept results consistent with a neural mechanism implicated in SA. Replication of these results within a larger sample is needed to allow for testing whether error-related MFC-sensory theta wPLI (associated with error monitoring) mediates the link between SA and memory biases for error events. Further, while the current results are suggestive of a neural mechanism by which errors are better encoded and subsequently remembered, it is important to further test if this proposed mechanism is predictive of the maintenance or worsening of SA via longitudinal methods. At shorter time scales, this could be tested by assessing whether changes in state SA are mediated, in serial, by enhanced error monitoring driving memory biases for error events. Similarly, the maintenance or worsening of trait SA could be assessed over the course of longer timescales (weeks/months). If subsequent work is able to provide more direct evidence in support of a neural mechanism implicated in SA, then this could inform the development of novel, brain-based treatment approaches, as it has already been demonstrated that MFC theta oscillations can be non-invasively manipulated [92], [93].

## Conclusions

In an effort to move beyond neural markers of “risk” and towards the identification of neural mechanisms implicated in SA, the current study provides evidence that error-related MFC theta oscillations (associated with error monitoring) impact what is encoded about social situations and subsequently remembered. Moreover, we demonstrate that SA is associated with enhanced error-related MFC theta oscillations and memory biases for error events. These findings introduce the possibility that error-related MFC theta oscillations could play a causal role in the etiology of SA. Nonetheless, the current results should be considered only as preliminary, proof-of-concept evidence for such a possibility, given the small sample and correlational nature of the current study. Future work should seek to replicate and extend these findings, employing longitudinal methods within larger and more diverse samples.

## Data Availability

The Psychopy task, questionnaires, data pre- and post-processing scripts, as well as data analyses scripts are publicly available on the following GitHub repositories: https://github.com/NDCLab/memory-for-error-mini, https://github.com/NDCLab/social-flanker-eeg-dataset. Deidentified data are available from the corresponding author upon request.

## Conflicts of Interest

The authors have no potential conflicts of interest to disclose.

## Funding Statement

Research reported in this publication was supported by the National Institute of Mental Health of the National Institutes of Health under award number R01MH131637 (Buzzell, Pettit), as well as through an FIU Center for Children and Families (CCF) Seed Funding grant (Hosseini).

## Acknowledgments

We would like to thank all undergraduate research assistants at the Neural Dynamics of Control Lab that assisted with data collection. We also thank the participants taking part in the study.

## Supplementary materials

Supplementary materials contain detailed steps of EEG data preprocessing and additional analyses to rule out an alternative interpretation explaining relations between SA and memory biases for error events.

## Notes

### Competing Interest Statement

The authors have declared no competing interest.

