## Supplementary materials for "Towards a mechanistic understanding of the role of error monitoring and memory in social anxiety"

### EEG preprocessing

Complete details of the EEG preprocessing stream are provided here, to supplement details provided in the main text. EEG data were preprocessed using MATLAB R2021b (MathWorks Inc., Sherborn, MA, USA), the EEGLAB toolbox, and a modified version of the MADE pipeline [1], [2]. Response offsets did not require correction, given the 1000 Hz sampling rate and mean offsets were measured to be ≤ 1 ms. Data were filtered via a high-pass filter at 0.1 Hz, followed by a low-pass filter with a passband of 49 Hz and a stopband of 59 Hz. The FASTER EEGLAB plugin [3] was used to identify/exclude bad channels. To further remove noise components originating from ocular and/or muscular activity, we performed independent component analysis (ICA). To improve ICA decomposition, a copy of each dataset was made, high-pass filtered at 1 Hz, and segmented into 1-second epochs [1]. If a channel had more than 20% artifacted epochs, that channel was removed from both the copied and original dataset [1]. No more than 10% of channels were identified as bad for all participants. ICA was then performed on the filtered/cleaned dataset. ICA weights were copied back to the original dataset and the adjusted-ADJUST algorithm [4], [5] used to identify independent components (ICs) corresponding to artifacts; identified ICs were subtracted from the data. Data were segmented into 3-second epochs (-1 to 2 seconds, relative to the response) and the DC-offset subtracted from each epoch. To eliminate any remaining ocular artifacts, epochs were rejected if a ±125 μV threshold was exceeded for channels located near the eyes. For all other channels not located near the eyes, if the ±125 μV threshold was exceeded, the channel was interpolated (unless more than 10% of channels in a given epoch exceeded the threshold, in which case the entire epoch was removed). Finally, a spherical spline interpolation was utilized to interpolate missing channels, after which channels were re-referenced to the average of all channels.

### Ruling out an alternative interpretation explaining relations between Social Anxiety (SA) and Memory Biases for Error Events

Within the main text, we report that SA is associated with memory biases for error events. While we interpret such memory biases as arising from enhanced error monitoring in high SA individuals, this cannot be confirmed based on behavioral data alone. This is because alternatively, high SA individuals could simply be more distracted by faces to begin with (preferentially attending to and encoding faces), which then causes errors to occur, as opposed to error monitoring driving the encoding of error events. However, as reported in the main text, our neural data present a pattern of results consistent with our hypothesis that memory biases for error events are driven by heightened error monitoring—not distraction by face stimuli. That is, SA symptoms were positively associated with heightened memory biases for error events as well as heightened error-related MFC theta oscillation patterns indicative of enhanced error monitoring. In particular, high SA individuals exhibited enhanced error-related MFC-sensory theta wPLI, which further predicted subsequent memory biases for error events. Here within the supplement, we present additional analyses of stimulus-evoked neural data from the Face-Flanker task, which fails to provide any evidence in favor of the alternative interpretation (that memory biases in SA are driven by distraction caused by face stimuli).

The alternative interpretation states that the relation between memory biases for error events and SA arises from individuals high in SA being more likely to be distracted by faces to begin with, preferentially attending to faces at stimulus onset (increasing the likelihood of encoding), which then causes error responses. Crucially, design of the Face Flanker task, which involves a stimulus onset asynchrony between the onset of face and target stimuli, as well as the recording of neural measures, allows for distinguishing between our original hypotheses and the alternative interpretation. If the alternative interpretation is correct, then neural responses to face onsets (which occur prior to target onsets) should be most pronounced on error (vs. correct) trials for high SA individuals and/or those exhibiting stronger memory biases for error events. Thus, in order to provide a direct test of this alternative interpretation, we extracted stimulus-locked neural responses to the onset of the face stimuli from the Face-Flanker task. Specifically, we extracted the N1/N170 event-related potential (ERP) component, which has been linked to face perception and is characterized by a negative deflection in the ERP waveform peaking ~170 ms after stimulus onset over lateral occipitotemporal scalp locations [6], [7], [8, p. 1]. To compute the N170 ERP component, mean amplitudes were separately extracted for error and correct trials within a 150-200 post-stimulus window within a bilateral cluster of occipito-parietal electrodes (22, 24, 53, and 55). Error-correct N170 ERP component difference scores were then computed.

The alternative interpretation makes two explicit predictions: 1) SA symptoms should be associated with an increase in the magnitude of the error-correct N170 difference score (negative correlation, given that the N1/N170 is a negative going voltage deflection); 2) memory biases for error events should also be associated with an increase in the magnitude of the error-correct N1/N170 difference score (negative correlation, given that the N1/N170 is a negative going voltage deflection). Each of these hypotheses were tested via two-tailed Pearson correlation tests. First, we found that SA symptom levels (assessed via SCAARED-social) were not significantly associated with the N1/N170 ERP component difference score, ($r(22) = 0.18$, $p = 0.39$). Moreover, we found no significant correlation between memory biases for error events and the N1/N170 ERP component difference score, ($r(22) = 0.09$, $p = 0.68$). Thus, neither of the predictions made by the alternative prediction were supported by the data. Taken together with the neural data reported in the main text, our results not only demonstrate that SA is associated with memory biases for error events, but also provide evidence that such memory biases may arise as the result of error-related MFC theta oscillations associated with error monitoring.
